# Temperature controlled high-throughput magnetic tweezers show striking difference in activation energies of replicating viral RNA-dependent RNA polymerases

**DOI:** 10.1101/2020.01.15.906032

**Authors:** M. Seifert, P. van Nies, F.S. Papini, J.J. Arnold, M.M. Poranen, C.E. Cameron, M. Depken, D. Dulin

## Abstract

RNA virus survival depends on efficient viral genome replication, which is performed by the viral RNA dependent RNA polymerase (RdRp). The recent development of high throughput magnetic tweezers has enabled the simultaneous observation of dozens of viral RdRp elongation traces on kilobases long templates, and this has shown that RdRp nucleotide addition kinetics is stochastically interrupted by rare pauses of 1-1000 s duration, of which the short-lived ones (1-10 s) are the temporal signature of a low fidelity catalytic pathway. We present a simple and precise temperature controlled system for magnetic tweezers to characterize the replication kinetics temperature dependence between 25°C and 45°C of RdRps from three RNA viruses, i.e. the double-stranded RNA bacteriophage Φ6, and the positive-sense single-stranded RNA poliovirus (PV) and human rhinovirus C (HRV-C). We found that Φ6 RdRp is largely temperature insensitive, while PV and HRV-C RdRps replication kinetics are activated by temperature. Furthermore, the activation energies we measured for PV RdRp catalytic state corroborate previous estimations from ensemble pre-steady state kinetic studies, further confirming the catalytic origin of the short pauses and their link to temperature independent RdRp fidelity. This work will enable future temperature controlled study of biomolecular complex at the single molecule level.

## Introduction

Genome replication is essential to any living organism. To achieve this task, every RNA virus encodes an RNA dependent RNA polymerase (RdRp) that synthesizes all the viral RNA, either to form messenger RNA for viral protein translation, or to produce new viral genomes that are enclosed into viral particles as they mature into infectious virions. Furthermore, the viral RdRp also evolves the viral genome either by incorporating mutations (1,2) while replicating the viral genome, or by assisting RNA recombination (3). The viral RdRp is a key player in successful infection, and is therefore the target of many antiviral drugs (4,5). Of special interest is the mechanochemical cycle of nucleotide incorporation occurring at the RdRp’s catalytic site (6). This site is conserved amongst RNA viruses (7), and resembles the one of the DNA and RNA polymerases in the A family with their typical cupped right hand shape (7-9). PV and Φ6 RdRps are model for the RdRps of positive-sense single-stranded (ss) and double-stranded (ds) viruses, respectively. Pre-steady-state kinetics, structural biology, molecular dynamics and next generation sequencing studies on PV RdRp and its genome have largely shaped our understanding on nucleotide selection and incorporation kinetics by viral RdRps (10), while structural and biochemical analyses on the Φ6 RdRp have provided insights into the de novo initiation mechanism (11-13). However, a complete description of the kinetic cycle of viral RdRp’s elongation is still lacking. Indeed, the ensemble techniques are unable to access RdRp kinetics on genome-long templates, i.e. kilobases (kb), and cannot interrogate rare asynchronous events, such as nucleotide misincorporation or antiviral nucleotide analogue incorporation in the presence of cognate canonical NTPs. Our recent single molecule studies on Φ6 and PV RdRps elongation kinetics have partly filled this gap, shedding light on the kinetic of elongation pauses of various biochemical origins (14-16). This work relied on the concomitant development of high throughput magnetic tweezers, a powerful single molecule force and torque spectroscopy technique enabling the characterization of protein-nucleic acids interactions at both high throughput (15,17-20) and high-resolution (21-23), and of a new analysis approach based on First-Passage statistics (24,25). However, the home-built magnetic tweezers instrument used for these studies did not include a temperature control system, and the experiments were therefore performed at room temperature. Ensemble kinetic assays have demonstrated the temperature dependence of RdRps activity, either in initiation for dengue and Zika virus RdRps (26), or in elongation for PV RdRp (27). Furthermore, temperature controlled experiments performed on *Escherichia coli* RNA polymerase (RNAP) at the single molecule level using optical tweezers have further informed on the mechanochemical cycle of nucleotide addition and on the nature of RNAP pauses, showing that off-pathway pauses have no enthalpic contribution, i.e. pause exit rate demonstrating no temperature dependence, whereas nucleotide addition rate is dominated by enthalpy, i.e. having strong temperature dependence (28,29). A temperature dependent study of viral RdRp’s elongation kinetics would thus significantly complement our current understanding of their mechanochemical cycle, and warrants the development of temperature controlled high throughput magnetic tweezers.

Recently, several studies have reported on the development of custom temperature controlled magnetic tweezers assays, i.e. relying either on a totally custom approach (home-built proportional-integral-derivative (PID) controller, heating elements and thermistors disposed at several locations on the microscope, and a custom graphic user interfaces (GUI)), or a dedicated commercial device (30-34). In the present study, we take advantage of a simple and robust commercial device from Thorlabs (originally designed to control the temperature on 1 inch diameter optical tubes and provided with its own GUI) to precisely control the temperature on an oil immersion microscope objective. Doing so, we are able to maintain the temperature within ∼0.1° from room temperature up to 60°C in the whole field of view, i.e. with dimensions of (0.5 × 0.4) mm, of a home-built high throughput magnetic tweezers.

We applied this temperature controlled high throughput magnetic tweezers to study the temperature dependence of the replication kinetics of three viral RdRps, i.e. Φ6, PV and HRV-C RdRps. On the one hand, we expect a different response to temperature for Φ6 and PV RdRps, as Φ6 and PV are respectively a plant pathogenic bacteria bacteriophage and a human virus, with respective host optimal temperature of 28°C and 37°C. On the other hand, we expect a similar response to temperature for PV and HRV-C RdRps, as both are human enteroviruses. Indeed, we show that PV and HRV-C RdRps present a steep and similar increase in nucleotide addition rate and in the exit rates for the short pauses (0.3-10 s), while the same kinetic states are lesser affected for Φ6 RdRp. Furthermore, we noticed that the long pauses (>20 s) probability is largely unaffected for all RdRp. For the three RNA virus we studied, we show that RNA virus replication rate is optimum near the optimum growth temperature of the host cell. Moreover, we believe that the easy implementation and low-cost of the temperature control system we have characterized makes it very attractive and will therefore be of large interest to the single molecule community using instruments that include an oil immersion objective, as well as for fluorescence microscopy and optical tweezers.

## Materials and Methods

### High throughput magnetic tweezers apparatus

The high throughput magnetic tweezers apparatus has been previously described (15,18,35). Shortly, it is a custom inverted microscope with a 50× oil immersion objective (CFI Plan Achro 50 XH, NA 0.9, Nikon, Germany), on top of which a flow cell is mounted. The assembly, surface preparation of the flow cell and nucleic acid tethering of the magnetic beads is described in the paragraph below. To apply an attractive force to the magnetic beads and stretch the nucleic acid tether, a pair of vertically aligned permanent magnets (5 mm cubes, SuperMagnete, Switzerland) separated by a 1 mm gap (36) are positioned above the objective; the vertical position and rotation of the beads are controlled by the M-126-PD1 and C-150 motors (Physik Instrumente PI, GmbH & Co. KG, Karlsruhe, Germany), respectively (**Figure 1A**). The field of view is illuminated through the magnets’ gap by a collimated LED-light source located above it, and is imaged onto a large chip CMOS camera (Dalsa Falcon2 FA-80-12M1H, Stemmer Imaging, Germany). The temperature control system is made of a flexible resistive foil heater with an integrated 10Ω thermistor (HT10K, Thorlabs) wrapped around an oil immersion objective (CFI Plan Achro 50 XH, NA 0.9, Nikon, Germany) and further insulated by several layers of kapton tape (KAP22-075, Thorlabs). The heating foil is connected to a PID temperature controller (TC200 PID controller, Thorlabs) to adjust the temperature within ∼0.1°C. We used the Thorlabs GUI to control the heating system via USB on the data acquisition computer.

**Figure 1:**
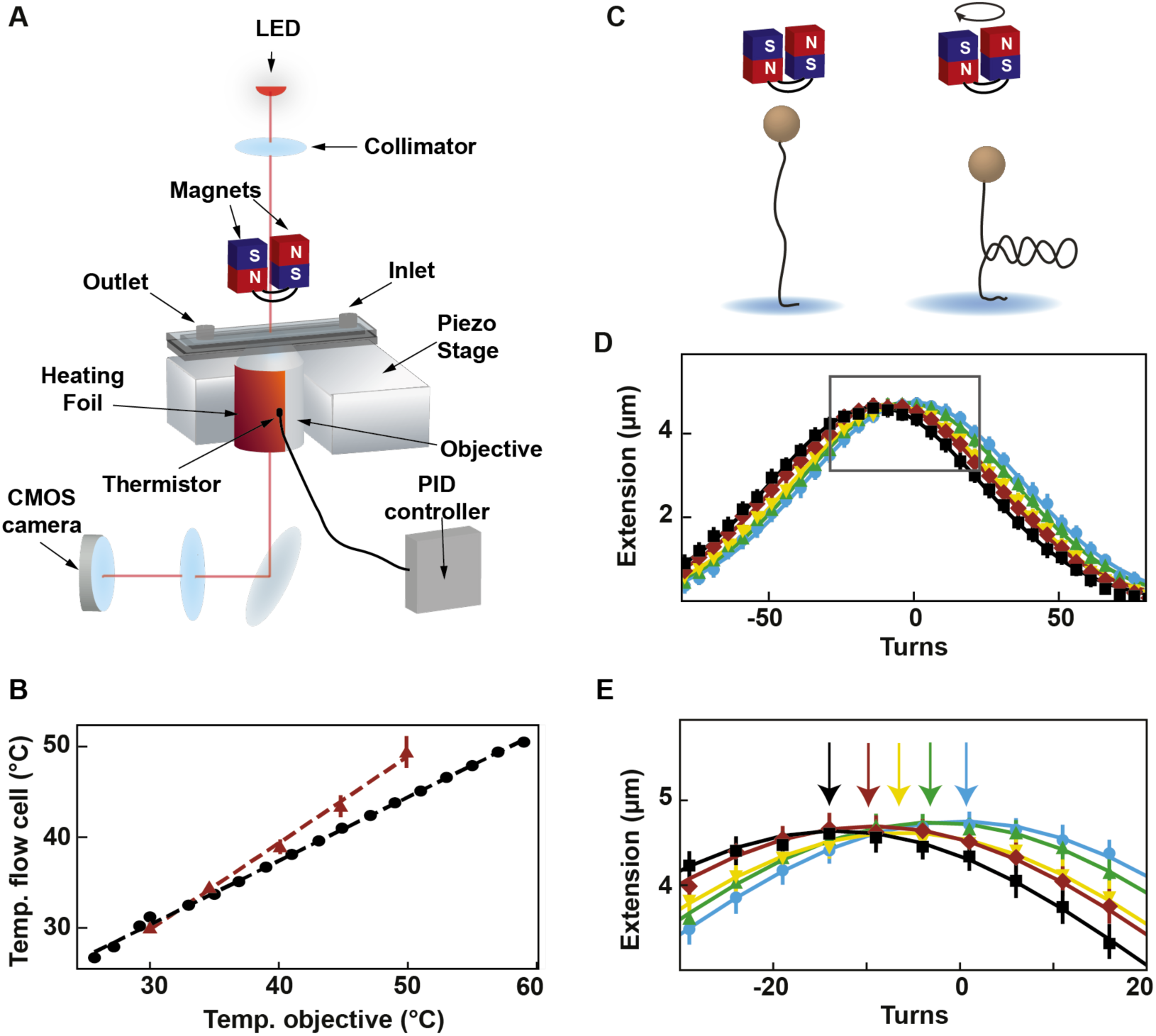
Description and calibration of the temperature control device installed in the high throughput magnetic tweezers assay. **(A)** Schematic of the magnetic tweezers assay. **(B)** Temperature measured at the surface of the flow cell as a function of the temperature measured at the objective by the thermistor of the heating foil. Red circles: temperature at the surface of the flow cell extracted from the extension-rotation experiments (**Material and methods**; error bars are the standard deviation for seven tethers). Black circles: temperature at the surface of the flow cell measured using a macroscopic thermistor (**Supplementary Figure 1B**). The dashed lines are linear fits. **(C)** Schematic of supercoiling of linear dsDNA. **(D)** The median rotation-extension of seven different MyOne magnetic beads tethered by 20.6 kbp dsDNA at either 30°C (light blue), 35°C (green), 40°C (yellow), 45°C (red) or 50°C (black), experiencing a 0.3 pN force. Each rotation-extension experiment for a given temperature is fitted by a Gaussian. **(E)** A magnification of the shift of the maximum extension position. Arrows indicate maxima.

### Flow cell fabrication

The flow cell assembly has been described previously (35). The flow cell was mounted on the magnetic tweezers setup and rinsed with 1 ml of 1×phosphate buffered saline (PBS). 100 µl of a 1:1000 dilution of 3 µm polystyrene beads (LB30, Sigma Aldrich, Germany) were added and after a 3-minute incubation, the flow cell was rinsed with 1 ml PBS. 40 µl of anti-digoxigenin Fab fragments (1 mg/ml) were added and the excess rinsed away with 1 ml PBS after 30 minutes incubation. The flow cell was then treated with bovine serum albumin (BSA, New England Biolabs) (100 mg/ml) for 10 minutes and rinsed with 1 ml PBS. To remove BSA excess from the surface, high salt buffer (10 mM Tris, 1mM EDTA pH 8.0, supplemented with 750 mM NaCl and 2 mM sodium azide) was flushed through the flow cell and rinsed 10 minutes later with TE buffer (TE supplemented with 150 mM NaCl and 2 mM sodium azide). To establish the temperature dependent rotation-extension experiments, we carefully mixed 10 µl of streptavidin coated MyOne magnetic beads (Thermofisher, Germany) and with ∼0.25 ng of 20.6 kbp long coilable double-stranded DNA tether, diluted it to 40 µl, and flushed the solution into the flow cell (35). For the RdRp experiment, we mixed 5 µl of dsRNA at ∼0.1 ng/µl with 20 µl of streptavidin coated M270 magnetic beads (Invitrogen), diluted it to 40 µl and flushed the solution into the flow cell. For both experiments, the flow cell was rinsed to remove unattached magnetic beads after few minutes of incubation.

### Coilable DNA construct for temperature calibration

The 20.6 kbp coilable DNA construct fabrication was described in Ref. (35).

### dsRNA construct

The construct employed here, which has been previously described in detail (15), a 4 kb long single-stranded splint to which is annealed four ssRNAs: a biotins-labeled strand to attach to the streptavidin-coated magnetic bead, a spacer, ∼2.9 kb template, and a digoxygenin-labeled strand to attach the magnetic bead to the surface glass surface. The template strand ends in 3’ with either a ssRNA flap made of 3 C residues followed by 15 U residues – used for Φ6 RdRp to catalyze de novo initiation, or a small hairpin with the sequence ACGCUUUCGCGT followed by 15 U residues to initiate PV and HRV-C RdRp catalyzed RNA synthesis via primer extension (27,37)(**Figure 2A**).

**Figure 2:**
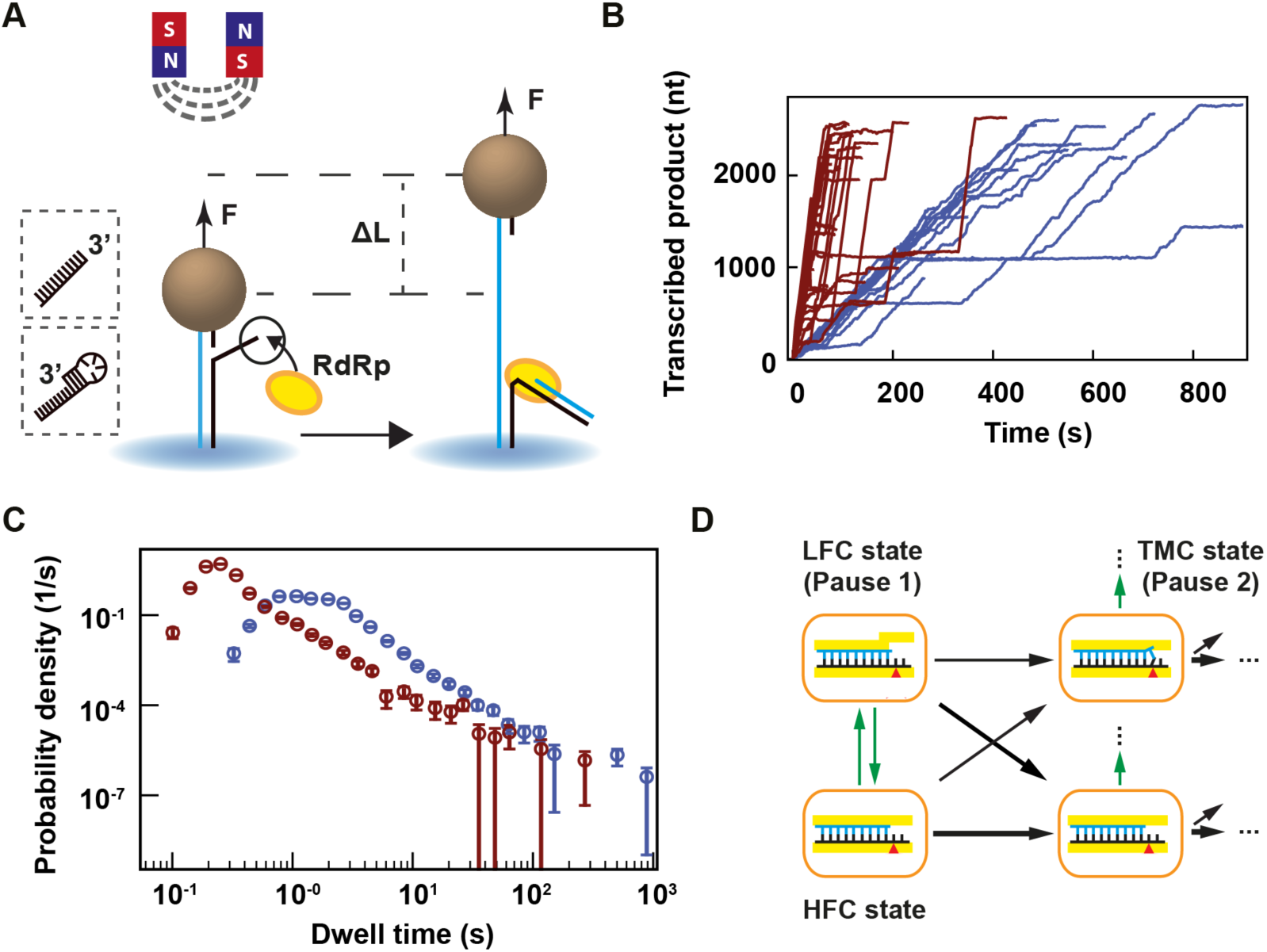
Temperature dependent magnetic tweezers assay show an increased replication rate for human rhinovirus C RdRp at increasing temperatures. **(A)** A schematic overview of the experimental assay (not to scale). A M-270 magnetic bead is tethered by a dsRNA that experiences a constant force F. The RdRp initiates at the 3’-end of the template strand, which ends either with a single-stranded 3’ overhang or a short hairpin (see insert on the left). In the presence of NTPs, the RdRp replicate and unwinds the template strand, creating a ssRNA tether with a difference in extension length ΔL in comparison to the dsRNA tether at a force F. **(B)** Low pass filtered (0.5 Hz) HRV-C RdRp activity traces at 25°C (blue) and 45°C (red), while applying a 30 pN constant force. **(C)** The dwell times probability density distribution extracted from at least 35 traces of HRV-C RdRp at either 25°C (blue) or 45°C (red) acquired as in (B). The error bars represent the 36% confidence interval obtained from 1000 bootstrapped datasets. **(D)** Kinetic model of nucleotide incorporation and misincorporation by RdRps (adapted from Ref. (15)). HFC: high fidelity catalytic state; LFC: low fidelity catalytic state; TMC: terminal mismatch catalytic state.

### Purification of Φ6 P2 RdRp

Recombinant N-terminally histidine-tagged Φ6 RdRp was expressed from plasmid pAA5 in *E. coli* BL21 (DE3) (38) and purified using Ni-NTA affinity column (Qiagen), HiTrap_TM_ Heparin HP column and HiTrap_TM_ Q HP column (GE Healthcare) as previously described (39). The purified protein was stored in 50% glycerol, 140 mM NaCl, 50 mM Tris-HCl pH 8.0, 0.1 mM EDTA, 0.1% Triton-X 100 at −20°C.

### Construction of HRV-C and PV RdRp bacterial expression plasmids

The HRV-C RdRp (3D gene) was cloned into the pSUMO bacterial expression plasmid using a similar procedure as described for PV RdRp (3D gene)(40). This system allows for the production of SUMO fusion proteins containing an amino-terminal hexahistidine tag fused to SUMO that can be purified by Ni-NTA chromatography and subsequently processed by the SUMO protease, Ulp1. Briefly, the HRV-C and PV RdRps coding region was amplified using respectively the HRV-C15 cDNA (Accession# GU219984.1) and the PV type 1 cDNA (Accession# V01149.1) as template, as described in Ref. (41). The PCR product of HRV-C RdRp was gel purified and cloned into the pSUMO plasmid using BsaI and SalI sites.

### Expression and purification of HRV-C and PV RdRps

HRV-C RdRp was expressed and purified using the same procedure reported for PV RdRp (40,42). Briefly, expression was performed at 25°C by auto-induction, cells harvested, lysed by French Press, subjected to PEI precipitation followed by AMS04 precipitation, Ni-NTA chromatography, cleavage by Ulp1, phosphocellulose chromatography, gel filtration and the protein concentrated using Vivaspin concentrators. *Reaction buffer for Φ6 P2*. The P2 reaction buffer is composed of 50 mM HEPES pH 7.9, 20 mM ammonium acetate, 3% w/v polyethylene glycol 4000, 0.1 mM EDTA (pH 8.0), 5 mM MgCl_2_, 2 mM MnCl_2_, 0.01% Triton X-100, 5% Superase RNase inhibitor (Life Technologies), 20 µg/ml BSA and 9 nM P2 RdRp.

### Reaction buffer for PV and HRV-C RdRps

The PV/HRV-C RdRp reaction buffer (coined PV reaction buffer) is composed of 50 mM HEPES pH 7.9, 5 mM MgCl_2_, 0.01% Triton X-100, 5% Superase RNase inhibitor (Life Technologies). To stall PV and HRV-C RdRps on the template, 500 nM of RdRp and 1 mM of ATP, CTP and GTP in PV reaction buffer were incubated in the flow chamber for ∼10 minutes, and the flow cell was subsequently rinsed.

### Single-molecule RdRp replication activity experiments

We select the dsRNA tether in the flow cell in reaction buffer as previously described (15). To prevent reinitiation on the template-product duplex when PV/HRV-C RdRp falls off, we first stalled the RdRp and rinsed the flow chamber with 400 µl of reaction buffer. Subsequently, we flushed in the reaction buffer, and grabbed the data for 30 minutes at 58 Hz camera acquisition rate, at a defined temperature and 30 pN force. The reaction buffer contains either 1 mM concentration of all NTPs for PV and HRV-C RdRps’ or 1 mM ATP/GTP and 0.2 mM CTP/UTP for Φ6 RdRp. The images are analyzed in real-time using custom-written routines in Labview 2017 and CUDA nVidia to extract the (x, y, z) position of up to ∼800 magnetic beads simultaneously (18). The micrometric change in tether extension upon RdRp activity (**Figure 3A**) is converted into numbers of transcribed nucleotides using the force-extension relationships for dsRNA and ssRNA constructs (15). RdRp activity traces are low-pass filtered at 0.5 Hz using a Kaiser-Bessel window, from which the dwell time were extracted as previously described (14,15,24).

**Figure 3:**
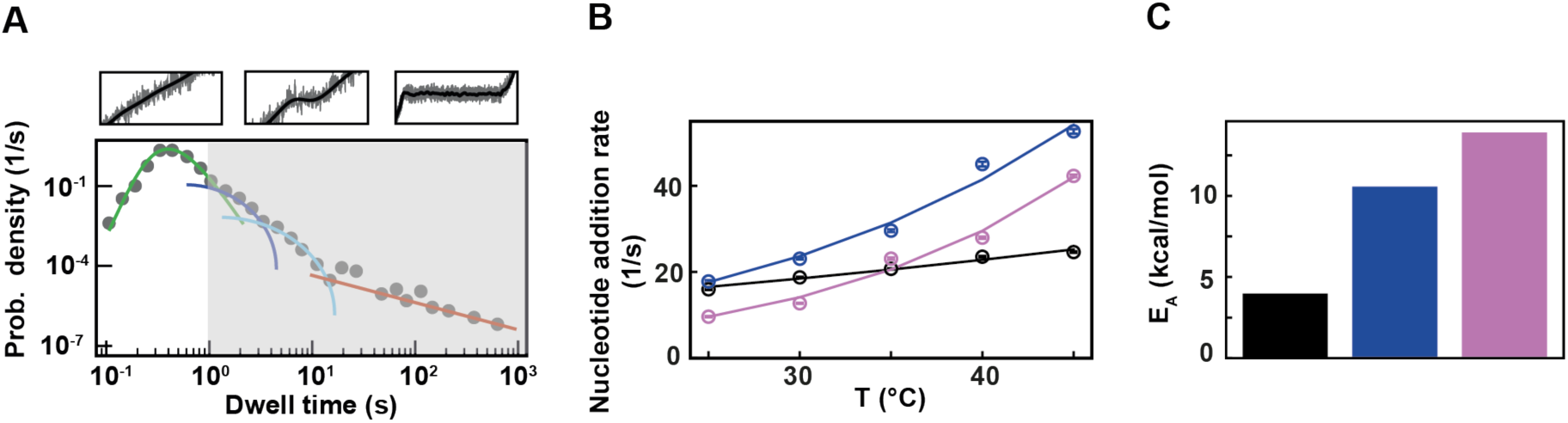
Φ6, PV and HRV-C RdRps show different nucleotide addition rates response to temperature increase. The corresponding data for each RdRp is represented in black, blue and pink, respectively in **(B)** and **(C). (A)** Section (not shaded) of the dwell time distribution that contributes to the MLE fit (green solid line) of the nucleotide addition (**Supplementary Figure 1D**). **(B)** The nucleotide addition rates (circles) as obtained from the maximum likelihood estimation (MLE) fits. The solid lines represent the respective fit of the Arrhenius equation. The error bars denote the standard deviation from 100 bootstraps of the MLE procedure (**Material and Methods**). **(C)** Activation energy E_A_ extracted from the fits in **(B)**.

### Stochastic-pausing model

There are many kinetic models that are consistent with the empirical dwell-time distributions we observe, and we here work under the assumption that the probability of pausing is low enough that there is only one rate-limiting pause in each dwell-time window. This assumption washes out most details of the kinetic scheme that connects pauses and nucleotide addition, but allows us to determine the general form of the dwell-time distribution without specifying how the pauses are connected to the nucleotide addition pathway

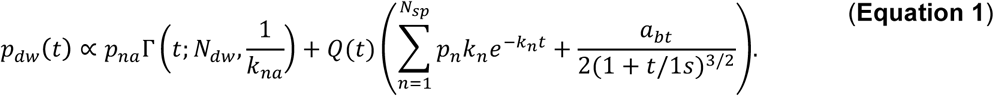

In the above expression, the gamma function in the first term contributes the portion *p*_*na*_ of dwell-times that originate in the RdRp crossing the dwell-time window of size *N*_*dw*_ base pairs without pausing; the second term is a sum of contributions originating in pause-dominated transitions, each contributing a fraction *p*_*n*_ of dwell-times; the third term captures the asymptotic power-law decay (amplitude *a*_bt_) of the probability of dwell-times dominated by a backtrack. The backtracked asymptotic term needs to be regularized for times shorter than the diffusive backtrack step. We have introduced a regularization at 1 s, but the precise timescale does not matter, as long as it is set within the region where the exponential pauses dominate over the backtrack. From left to right, each term of **Equation 1** is dominating the distribution for successively longer dwell-times.

A cut-off factor *Q* (*t*) for short times is introduced to account for the fact that the dwell-time window includes *N*_*dw*_ nucleotide-addition steps,

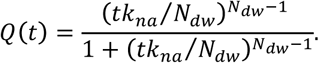

The fit results dependence on these cut-offs is negligible as long as they are introduced in regions where the corresponding term is sub-dominant. Here the cut is placed under the center of the elongation peak, guaranteeing that it is placed where pausing is sub-dominant.

### Maximum likelihood estimation

The normalized version of **Equation 1** is the dwell-time distribution fit to the experimentally collected dwell-times {*t*_*i*_}_*i*_ by minimizing the likelihood function (43)

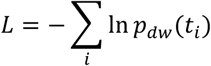

with respect to rates and probabilistic weights. We calculate the errors in our parameter estimates by bootstrapping (44) the system 100 times, and report the one-sigma confidence intervals among the bootstrapped data sets.

### Dominating in a dwell-time window vs. dominating in one step

The fractions *p*_*n*_ represent the probability that a particular rate *k*_*n*_ dominates the dwell-time. We want to relate this to the probability *P*_*n*_ that a specific exit rate dominates within a one-nt transcription window. Assuming we have labelled the pauses so that *k*_*n*-1_ > *k*_*n*_, we can relate the probability of having rate *n* dominating in *N*_*dw*_ steps to the probability of having it dominate in one step through

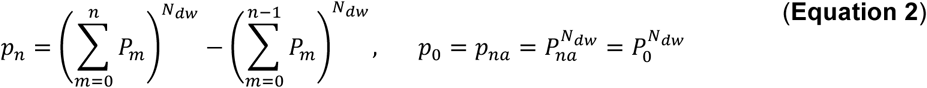

The first term in **Equation 2** represents the probability of having no pauses longer than the *n*^*th*^ pause in the dwell-time window, and the second term represents the probability of having no pauses longer than the (*n* - 1)^*th*^ pause. The difference between the two terms is the probability that the *n*^*th*^ pause will dominate. This can be inverted to yield a relation between the single-step probabilities (*P*_*n*_) and the dwell-time window probabilities (*p*_*n*_)

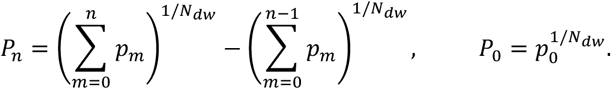

This relationship has been used throughout the manuscript to relate our fits over a dwell-time window to the single-step probabilities.

## Results

### Establishing temperature control of high throughput magnetic tweezers

To perform a temperature-controlled experiment in a magnetic tweezers assay, one needs to maintain the temperature setpoint constant in the field of view. To fulfill this condition using a magnetic tweezers instrument, it is sufficient to only control the temperature at the objective (given that an oil immersion objective is used) (30). Therefore, we have mounted a temperature control device on the objective that is derived from Thorlabs parts, meaning a flexible resistive foil heater including a thermistor that is strapped around the oil immersion objective (**Supplementary Figure 1A**) and a PID controller to adjust the temperature setpoint (**Figure 1A, Materials and Methods**). This system has several advantages in comparison with custom ones: it is commercially available, low price, and it comes with a graphic user interface (GUI) to adjust the temperature from the data acquisition computer. We first calibrated the temperature measured on the objective side versus the temperature measured in the flow cell. To this end, we used a digital portable thermometer (Gochange, Amazon.de, ±1.5 % error), which ∼1 mm^3^ temperature probe was inserted in the flow cell and immerged in ultrapure water (**Supplementary Figure 1B**). In absence of heating, i.e. at room temperature, the two sensors reported the same temperature. We repeated this measurement several times at different temperature setpoints assigned on the GUI to the PID controller, from 25°C to 60°C, which is the maximum temperature tolerated by the objective. We observed a linear relationship between the temperature at the objective and the temperature in the flow chamber, i.e. *T*_*flow cell*_ (*T*_*obj*_) = (0.7 * *T*_*obj*_ + 9.23) °*C* (**Figure 1B**). Because the probe integrates the temperature over a cubic millimeter volume, the thermometer measurement may not be representative of the temperature within few microns above the bottom coverslip surface of the flow cell. To perform an in situ calibration directly at the surface, we took advantage of the well-characterized temperature dependence of the DNA helical twist, which decreases by ∼11°/°C/kbp (45-47). Magnetic tweezers are an ideal technique to measure the twist of a coilable molecule, such as fully double-stranded DNA, as they enable a precise control of the torque applied on such molecule (**Figure 1C**). We performed several DNA rotation-extension experiments at 0.3 pN and at different temperatures using a coilable DNA molecule (30,48) (**Figure 1D, Materials and Methods**). The extension of DNA molecules at different supercoiling density shows a typical bell-like shape distribution, with the maximum extension centered at zero turn (**Figure 1D**). This shape is well fitted by a Gaussian function, which enables the evaluation of the zero turn maximum extension (**Figure 1E**). If DNA helical twist decreases, the zero turn position shifts towards the negative turns. If DNA helical twist increases, the zero turn position now shifts towards the positive turns. Changing the temperature by steps of 5°C from 30°C to 50°C, and centering at zero turn the maximum extension obtained at 30°C, we observed that the rotation-extension traces shift towards the negative turns upon temperature increases, as expected from a reduction in DNA helical twist (**Figure 1DE**). Converting the shift of the peak of the traces into temperature variation using the above described relationship between DNA helical twist and temperature, we extracted almost an one-to-one linear relationship between the temperature measured at the surface of the flow chamber and on the side of the objective, i.e. *T*_*flow cell*_ (*T*_*obj*_) = (0.95 * *T*_*obj*_ + 1.18) °*C* (**Figure 1B**). As our magnetic tweezers assay has a very large field of view of ∼(0.5×0.4) mm, we evaluated the homogeneity of the temperature over the whole field of view. To this end, we performed a rotation extension experiment at both at 25°C and 45°C with the same DNA tether at five different locations in the field of view by moving the flow cell with the micrometric screws of the stage, and we did not measure any difference of the rotation-extension traces for a given temperature. This result confirms that the temperature is constant over the whole field of view (**Supplementary Figure 1C**) (32). We have now shown that using a simple and commercially available temperature control system, we are able to adjust and maintain the temperature in the entire field of view of a high throughput magnetic tweezers instrument. This assay is thus suitable to investigate the temperature dependence of viral RdRps elongation kinetics.

### Setting up high throughput magnetic tweezers to study viral replication

We have recently developed a high throughput magnetic tweezers assay to investigate viral RdRp replication kinetics (14-16). To this end, a dsRNA construct tethers a magnetic bead to the glass coverslip surface of the flow cell (**Figure 2A, Materials and Methods**). The 3’-end of the template RNA is terminated by either a blunt ssRNA to initiate Φ6 RdRp (49), or by a small hairpin to initiate PV or HRV-C RdRps (14,37) (**Figure 2A**). Following successful initiation, the RdRp displaces the template strand from the tethering strand, converting the tether from dsRNA to ssRNA, which increases linearly the end-to-end extension of the tether by ΔL at constant force F and is subsequently converted from microns to nucleotides (15) (**Figure 2A**). Using high throughput magnetic tweezers, dozens of RdRp activity traces are simultaneously acquired (**Figure 2B, Supplementary Figure 2AC**). In this work, we study the influence of temperature on the replication kinetics of three model RdRps.

The direct observation of HRV-C RdRp traces show that an increase in temperature from 25°C to 45°C dramatically increases its average replication rate (**Figure 2B**). This is also observable for PV RdRp, and, to a smaller extent, for Φ6 RdRp (**Supplementary Figure 2AC**). To further quantitate these observations, we performed a dwell time analysis of the 0.5 Hz low pass filtered traces. Specifically, the traces were scanned with non-overlapping windows of 10 nt to evaluate the duration of ten successive nucleotide incorporations, after which the collected dwell times are assembled into a probability density distribution (**Figure 2C, Supplementary Figure 2BD, Supplementary Figure 3**) (14,15,24). The dwell time analysis is a direct measure of the event(s) that dominated in time over ten successive nucleotide incorporations, i.e. either the incorporation of the ten nucleotides only or a pause that interrupted the ten nucleotides addition (**Supplementary Figure 1D**). Looking at the dwell times distribution extracted from the HRV-C traces acquired at 45°C in a log-log graph (**Figure 2B**), the bins describe a bell-like shape until ∼0.4 s, then a shoulder from ∼0.4 s till ∼3 s, and finally a straight line for the longer dwell times (**Figure 2C, Supplementary Figure 1D**). Each region of the distribution is the temporal signature of a specific biochemical event in elongating RdRps. To describe these events, we previously introduced four probability distribution functions: to fit the bell-like shape region, a gamma distribution that describes ten successive nucleotide additions cycles without pause; to fit the shoulders, two exponentially distributed pauses (Pause 1 and Pause 2); and finally to fit the dwell times dominated by the longest pauses, a power law distribution (15,16,50) (**Supplementary Figure 1D, Material and Methods**). We have previously showed (14,15) that RdRp incorporates cognate NTPs fast through the high fidelity catalytic (HFC) state (**Figure 2D**, gamma distribution), and rarely interconverts into another conformation, i.e. the low fidelity catalytic (LFC) state (**Figure 2D**), from which the RdRp slowly incorporates either a cognate NTP appearing as Pause 1 (diagonal arrow in **Figure 2D**) or a non-cognate NTP. In the latter, the nucleotide addition following the mismatch addition is done even slower than Pause 1 through a state we coined the terminal mismatch catalytic (TMC) state (**Figure 2D**) and appears in the traces as Pause 2. Furthermore, we also showed that the power law distributed pause relates to RdRp backtracking (16,50). Using a maximum likelihood estimation (MLE) procedure, we extracted the parameters for each distribution, i.e. exit rates and probabilities (**Material and Methods, Supplementary Table 2, Supplementary Figure 3**) (24). Having acquired and analyzed viral replication traces at several temperatures, we now describe how the parameters extracted from the MLE procedure vary for Φ6, PV and HRV-C RdRps, at constant force, i.e. 30 pN, and saturating NTPs concentration, i.e. 1 mM.

### The nucleotide addition rates of RdRps from an environmental phage and from two human viruses show different responses to temperature increase

As noted above, HRV-C and poliovirus RdRp show a dramatic increase in average elongation rate when increasing the temperature (**Figure 2BC, Supplementary Figure 2CD**), while only a mild effect is observable for Φ6 RdRp (**Supplementary Figure 2AB**). We first look into the pause-free nucleotide addition rate described by the bell-like shape gamma distribution (**Figure 3A, Supplementary Figure 1D**). For temperature varying from 25°C to 45°C, the nucleotide addition rate of PV RdRp increases from (17.8 ± 0.3) nt/s to (52.7 ± 0.5) nt/s, i.e. almost a 3-fold increase, and in agreement with previous bulk measurements (27). Similarly, HRV-C RdRp nucleotide addition rate strongly increases, i.e. from (9.6 ± 0.1) nt/s to (42.3 ± 0.3) nt/s, i.e. a 4-fold increase. On the other hand, Φ6 RdRp nucleotide addition rate only varies from (16.0 ± 0.1) nt/s to (24.6 ± 0.2) nt/s for the same temperature span, in line with a previous ensemble measurement (51) (**Figure 3A, Supplementary Table 2**). Investigating the temperature response of a reaction, one is able to determine whether the reaction is exothermic or endothermic from the Van’t Hoff equation by determining whether the sign of the reaction rate derivative by the temperature is negative or positive, respectively. Furthermore, it enables the measurement of the activation energy *E*_*A*_ of the reaction by fitting the reaction rate, i.e. nucleotide addition or pause exit, as a function of the temperature with the Arrhenius equation:

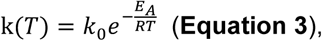

where *k* is the rate constant, *k*_0_ is the frequency factor, *T* is the absolute temperature, and *R* is the universal gas constant (52) (**Figure 3B**). As the nucleotide addition rate increases with temperature, the reaction is endothermic. Evaluating the activation energy (± standard deviation) of the nucleotide addition rate for the three RdRps using **Equation 3**, we found *E*_*A*, Φ6_ = (4.1 ± 0.4)*kcal/mol, E*_*A,PV*_ = (10.7 ± 1.0)*kcal/mol* and *E*_*A,HRV*-*C*_ = (14.3 ± 1.2) *kcal/mol* (**Figure 3C**). Φ6 and PV RdRp nucleotide addition rates are force insensitive (14,15), which means that the nucleotide addition rate is not limited by the polymerase ratchet-like translocation to the n+1 position. Therefore, the activation energy relates to another rate limiting process, such as the phosphoryl transfer (6). Conclusively, our results confirm a previous pre-steady state kinetic estimation of the free energy landscape of nucleotide addition for PV RdRp (6). Our results demonstrate the large impact of temperature on the nucleotide addition rate for human viruses, i.e. PV and HRV-C RdRp, and the mild impact on the environmental virus, i.e. the RdRp of *Pseudomonas* bacteriophage Φ6.

### Human virus and environmental bacteriophage RdRps elongation competent pauses are differently affected by temperature

We then investigated how Pause 1 and Pause 2 kinetics and probabilities (**Figure 4A** and **E**) are affected by temperature. We previously showed that Pause 1 and Pause 2 are the signature of slow nucleotide addition cycles that appear as short pauses in the RdRp elongation activity traces (14,15). For PV and HRV-C RdRps, Pause 1 exit rate increases dramatically, i.e. by 3 and 6-fold, respectively (**Supplementary Table 2, Figure 4BF**), which is further highlighted by their respective large Pause 1 activation energy, i.e. *E*_*A,PV*_ = (11.0 ± 1.1)*kcal/mol* and *E*_*A,HRV*-*C*_ = (15.2 ± 4.9)*kcal/mol* (**Supplementary Table 2, Figure 4BC**), similar to the nucleotide addition activation energy (**Figure 3C**). Pause 2 exit rate also experiences a steep increase, i.e. by 4- and 10-fold for PV and HRV-C RdRp’s, respectively, and eventually saturates at 35°C (**Supplementary Table 2, Figure 4F**). Of note, to achieve reasonable fits to the HRV-C dwell time distribution collected at 40°C, we were forced to restrict Pause 2 exit rate *k*_2_ > 0.2 *s*^-1^ (**Supplementary Figure 3**). The large increase of Pause 2 exit rates is further supported by an activation energy larger than for Pause 1, i.e. *E*_*A,PV*_ = (23.2 ± 0.5)*kcal/mol* and *E*_*A,HRV*-*C*_ = (34.3 ± 1.8)*kcal/mol* (**Equation 3, Figure 4G)**. Interestingly, Pause 1 and Pause 2 exit rates for PV RdRp increase with force (14), which suggests that forward translocation is the rate limiting step of slow nucleotide addition through this parallel kinetic pathway (**Figure 2D**). The large activation energy of Pause 2 is therefore consistent with the barrier to RdRp translocation induced by a nucleotide mismatch and is in agreement with previous ensemble measurements (6). Looking now at Φ6 RdRp, we observed a very different trend. Indeed, Pause 1 and Pause 2 exit rates are relatively stable, i.e. *k*_1_ = (1.7 ± 0.4) *s*^-1^ and *k*_2_ = (0.26 ± 0.09) *s*^-1^ respectively (mean ± standard deviation, **Supplementary Table 2, Figure 4BF**). Therefore, the Arrhenius equation fits poorly to Φ6 RdRp pause exit rates evolution with temperature (**Figure 4BF**), leading to a poor estimation of the activation energies of Pause 1 and Pause 2 (**Figure 4CG, Supplementary Table 2**). Off-pathway pause exit rates have been shown to be temperature insensitive in *E. coli* RNAP (28,29), and could offer an attractive explanation for the relative insensitivity of Pause 1 and Pause 2 from Φ6 RdRp. However, Φ6 RdRp elongation rate also increases little with temperature, and therefore we suspect that rate-limiting step in nucleotide addition is not thermally activated in the temperature range we explored. Pause 1 probabilities are largely unaffected by temperature for Φ6 and PV RdRps, while it decreases when temperature increases for HRV-C (**Figure 4D**). However, the nucleotide addition rate and Pause 1 HRV-C dwell time distributions have a strong overlap at low temperature (**Figure 3B, Supplementary Figure 3**), which may bias the probability we measured. Pause 2 probability is largely constant for all RdRps (**Figure 4H**). As for nucleotide addition rate, we observe a very different response to temperature for the human RNA virus RdRps and the environmental dsRNA bacteriophage, which could be an evolutionary advantage for a bacteriophage to not respond too strongly on natural environmental temperature variation in the room temperature range.

**Figure 4:**
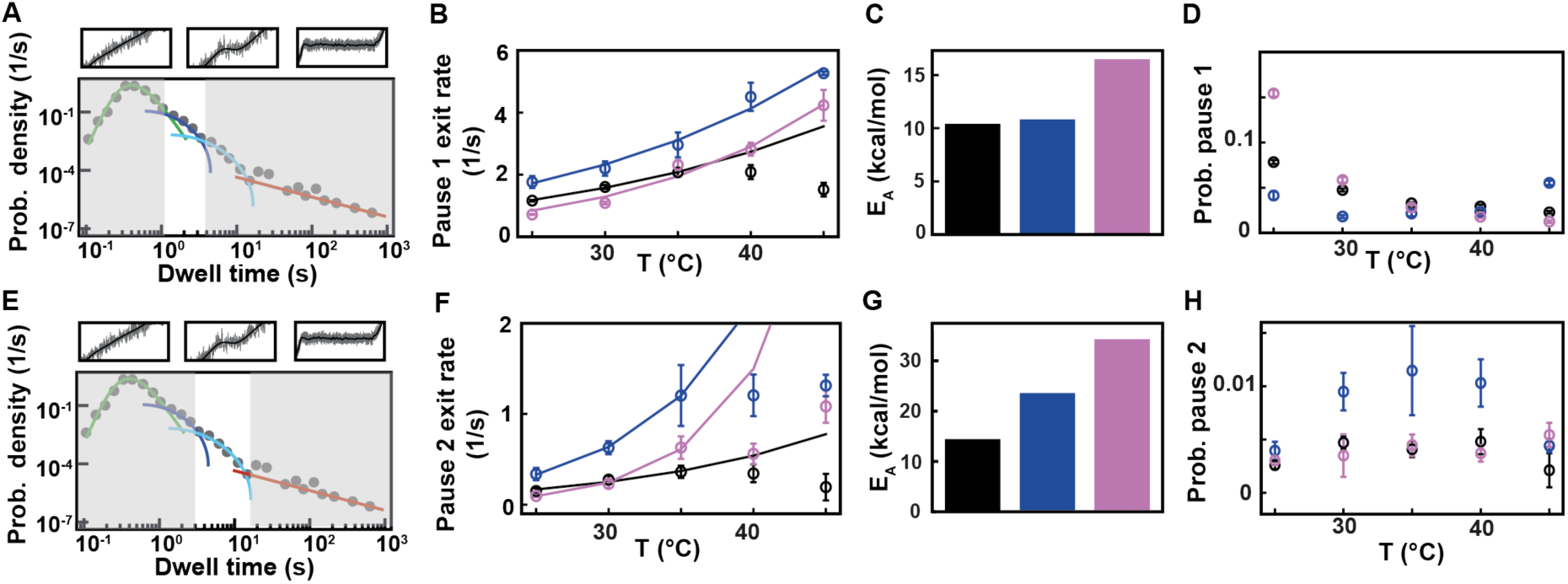
Pause 1 and Pause 2 exit rates and probabilities of Φ6, PV and HRV-C RdRps demonstrate different activation by temperature. The data for each RdRp is represented in black, blue and pink, respectively. **(A)** Section (not shaded) of the dwell time distribution that contributes to the MLE fit (dark blue solid line) of Pause 1. **(B)** The pause 1 exit rates (circles) extracted from the MLE fits as a function of the temperature. The lines represent the fitted Arrhenius equation. For Φ6 RdRp, only the data at 25°C, 30°C and 35°C were considered for the fit. **(C)** Activation energy for Pause 1 exit rate. **(D)** The probabilities to be in Pause 1 state as a function of the temperature. **(E)** Section (not shaded) of the dwell time distribution that contributes to the MLE fit (light blue solid line) of Pause 2. **(F)** The Pause 2 exit rates (circles) extracted from MLE fits. The lines represent the fitted Arrhenius equation. Only the data at 25°C, 30°C and 35°C were considered for the fit. **(G)** Activation energy for Pause 2 exit rate. **(H)** The probabilities to be in Pause 2 state as a function of the temperature. Error bars in B, D F and H denote the standard deviation extracted from 100 bootstraps of the MLE procedure.

### Backtrack related long pause probability mildly decreases with temperature increase for Φ6 RdRp, while remaining constant for PV and HRV-C RdRps

We next look at the last part of the dwell time distribution, which describes the pauses longer larger than 20 s (**Figure 5A**) and are related to polymerase backtrack, i.e. a backward diffusion of the polymerase on its template leading to an RNA product 3’-end out of register. The polymerase eventually returns in register by hopping from base to base with a forward and backward rate. This is modeled by a one dimensional diffusion process through a periodical energy landscape, and is described in the dwell time distribution by a power law distribution with two parameters, i.e. a −3/2 exponent and a characteristic hopping rate (28,50,53) (**Material and Methods**). While we have previously observed Φ6 RdRp backtracking (16), this behavior had remained elusive for PV RdRp in absence of the nucleotide analogue T-1106 (14). We suspected that the absence of long backtrack pauses in our previous study likely originated from an RNase contamination of the PV RdRp stock, which consequently decreased the tether lifetime and prevented the observation of these rare and long pauses. Having improved PV and HRV-C RdRp purification protocols (**Material and Methods**), we are now able to characterize these backtrack pauses for these two enterovirus RdRps (**Figure 5, Supplementary Figure 2C**). The average hoping rate is hidden in the dwell time distribution behind the shoulder of Pause 1 and Pause 2, as they statistically dominate, and we have therefore decided to fix this parameter at 1 s^-1^, a value that has been directly measured for *E. coli* RNAP (50,54). Using this fixed hoping rate, we evaluated the probability for the polymerase to enter a backtrack pause as a function of the temperature. For temperatures varying from 25°C to 45°C, Φ6 RdRp probability to enter a backtrack pause mildly decreases, i.e. from 0.0027 to 0.0008, while for PV and HRV-C RdRps, this probability is largely constant, i.e. 0.0008 ± 0.0002 and 0.0009 ± 0.0002 (mean ± standard deviation), respectively (**Figure 5B, Supplementary Table 2**). Though we should be cautious with the extracted probabilities as we do not know the average hopping rate of a backtracking RdRp, we observe a different backtracking behavior between an environmental bacteriophage RdRp and human virus RdRps.

**Figure 5:**
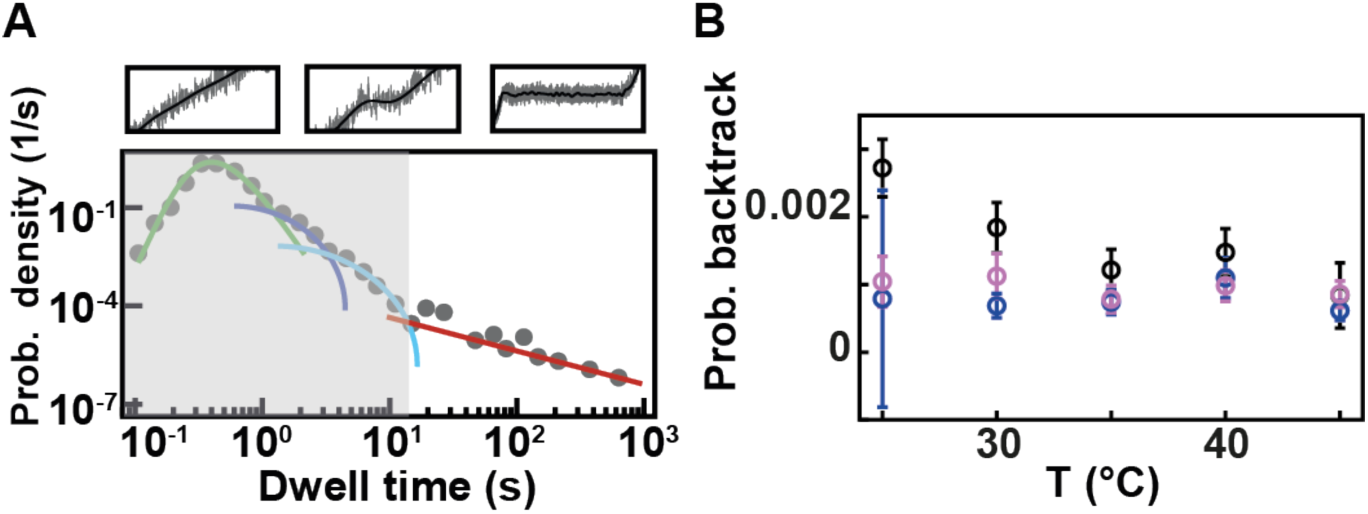
Temperature has little effect on backtrack probabilities for Φ6, PV and HRV-C RdRps. **(A)** Section (not shaded) of the dwell time distribution that contributes to the fit (red solid line) of the backtrack pause. **(B)** The probabilities for backtrack (circles) as obtained from MLE fits. The error bars denote the standard deviation extracted from 100 bootstraps of the MLE procedure. The data for Φ6, PV and HRV-C RdRps is represented in black, blue and pink, respectively.

## Discussion

Single molecule biophysics has revolutionized our view of molecular biology by enabling the observation of the kinetic processes of single enzymes, increasing significantly our understanding of these complex molecular motors (55). In particular, single molecule biophysics has further complemented the existing wealth of knowledge collected in ensemble measurements with transient, rare and asynchronous kinetic events to assemble complete mechanochemical pathways. However, while bulk studies are easily performed at physiological temperature, i.e. 37°C for human and *E. coli* enzymes, many single molecule studies are performed at room temperature (∼21°C) because of the technical difficulties at inserting a temperature control system within home-built high-end microscopes. Here, we present a simple temperature control system with several advantages: low cost, commercially available (Thorlabs), and easily mounted on an oil immersion objective. We calibrated the temperature in the field of view of a home-built magnetic tweezers instrument by performing rotation-extension experiments on a supercoiled DNA at several temperatures, using the well-known relationship describing the decrease in DNA helical twist as a function of temperature and achieving better performance than using a macroscopic thermometer probe.

We have previously investigated the kinetics of PV RdRp at room temperature, extracting a slow nucleotide addition rate of ∼10 nt/s, even at saturating concentration of NTPs and high assisting force (14). This result was consistent with previous bulk measurements performed at the same temperature on the same enzyme (27). Having established a temperature controlled magnetic tweezers assay, we have presented in this study how temperature affects the replication kinetics of RdRps from several different viruses, i.e. Φ6 (a bacteriophage infecting a plant pathogenic bacteria), PV and HRV-C (two human enteroviruses). These RdRps responded very differently to temperature change. Φ6 RdRp is only mildly affected in the temperature range we probed here, which is in line with previous work (51). *Pseudomonas syringae*, Φ6 natural host, grows at wide range of environmental temperatures (from 0 up to 36°C) with an wide optimum around ∼28°C (56). However, the reproduction cycle of Φ6 is restricted in temperatures above 30°C (57). Consequently, we suggest that Φ6 RdRp has evolved to support viral genome replication under different environmental temperatures, and therefore is not thermally activated above 25°C. On the other hand, the kinetics of PV and HRV-C RdRps are strongly affected by the increase in temperature. The data we present here definitely supports that the temperature is the predominant factor behind the slow replication we previously observed, as we have now measured nucleotide addition rates similar to the one measured in bulk at the same temperatures (27). Interestingly, PV and HRV-C RdRp showed a parallel trend, with PV nucleotide addition rate saturating at a lower temperature, suggesting that PV thrives at a slightly lower temperature than HRV-C, which may favor the infection of specific organs or cell-type with different optimal temperature. Interestingly, the activation energy we measured for the nucleotide addition rate and Pause 1, and the one for Pause 2 are both consistent with the energy barrier for cognate and non-cognate nucleotide addition, respectively, estimated by pre-steady state kinetic analysis (6). This result further supports our model where Pause 1 is a slow nucleotide addition event and Pause 2 is the signature of mismatch nucleotide addition (14,15)(**Figure 2D**), and suggests that temperature does not affect nucleotide misincorporation rate (Pause 2 probability is largely constant). Finally, we also show here that the replication traces of non-de novo initiating RdRps, i.e. PV and HRV-C, present long pauses related to polymerase backtrack, similarly to de novo initiating Φ6 RdRp (**Figure 5B**) (16). We anticipate that this behavior is ubiquitous in all RNA virus RdRps and may be an important feature for viral replication. On a technical note, we show here that our analysis framework is particularly suitable to detect even very short lived pauses, i.e. <1 s, while standard pause-removal approaches can only efficiently detect pauses longer than 1 s (58). Therefore, we are confident that our approach enables a good and less biased estimation of the nucleotide addition rate.

In conclusion, we have established a simple and easy to set up objective-based temperature control system for magnetic tweezers. This approach is easily applicable to any microscope using an immersion objective to ensure a good heat transfer. We believe that the simplicity and availability of our assay makes it very attractive, and will therefore be widely applied in the field. In addition, the RdRps replication kinetics study brings new evidences in support of our model of nucleotide mismatch addition.

## Supporting information

Supplementary Information for Seifert et al.

## Author Contributions

DD designed the research. MS and DD designed and performed the single molecule experiments. JJA, MP and CEC provided the recombinants RdRps. FSP made the RNA construct for the single molecule experiments; MS and PvN wrote the custom analysis software; MS analyzed the data; MD performed the MLE fits on all the data sets; all the authors discussed the data; MS and DD wrote the article and revised it based on the comments from the other authors.

## Acknowledgement

DD was supported by the Interdisciplinary Center for Clinical Research (IZKF) at the University Hospital of the University of Erlangen-Nuremberg. JJA and CEC are supported by grant AI045818 from NIAID, NIH. MMP was funded by Sigrid Juselius Foundation and Jane and Aatos Erkko Foundation. The use of the facilities and expertise of the Instruct-HiLIFE Biocomplex unit, a member of Biocenter Finland, Instruct-FI and Instruct-ERIC, is gratefully acknowledged. Authors thank Riitta Tarkiainen for excellent technical assistance.

