## Supplementary Information for Seifert et al. for "Temperature controlled high-throughput magnetic tweezers show striking difference in activation energies of replicating viral RNA-dependent RNA polymerases"

**Supplementary Table 1** The median shift of the rotational zero and the median temperature in the flow call as calculated using equation $\Delta Tw(T) = (-11.5 \pm 1.0)°/(°C\cdot kbp)$ from Ref. (1).

| Temperature objective (°C) | Median ± std rotational zero shift (turns) | Median ± std temperature flow cell (°C) |
|---|---|---|
| 30 | 0 ± 0 | 30 ± 0 |
| 34.6 | 3 ± 0.78 | 34.5 ± 0.4 |
| 40.1 | 6 ± 1.75 | 39.0 ± 0.8 |
| 44.8 | 9 ± 1.55 | 43.4 ± 1.2 |
| 49.9 | 13 ± 1.67 | 49.4 ± 1.8 |

**Supplementary Table 2. Kinetic parameters of the present study.** Nucleotide addition rates, Pause 1 and Pause 2 exit rates, their respective probabilities, and backtrack probability of Φ6, poliovirus and HRV-C RdRps obtained from the MLE fit of the model described in **Material and Methods**. The activation energy $E_A$ extracted from a fit of the Arrhenius equation to the rates. The blue lines denote the temperature range used to fit the Arrhenius equation.

| | nucleotide addition rate (1/s) ± std* | pause 1 exit rate (1/s) ± std | pause 2 exit rate (1/s) ± std | probability pause 1 ± std | probability pause 2 ± std | probability backtrack ± std | Ea kcat (kcal/mol) ± std | Ea k1 (kcal/mol) ± std | Ea k2 (kcal/mol) ± std |
|---|---|---|---|---|---|---|---|---|---|
| **Φ6** | | | | | | | | | |
| **25°C** | 16.0 ± 0.1 | 1.16 ± 0.02 | 0.15 ± 0.03 | 0.078 ± 0.002 | 0.0026 ± 0.0004 | 0.0027 ± 0.0004 | | | |
| **30°C** | 18.7 ± 0.1 | 1.60 ± 0.06 | 0.28 ± 0.04 | 0.047 ± 0.002 | 0.0047 ± 0.0006 | 0.0018 ± 0.0004 | | | |
| **35°C** | 20.7 ± 0.1 | 2.07 ± 0.12 | 0.36 ± 0.07 | 0.033 ± 0.001 | 0.0041 ± 0.0007 | 0.0012 ± 0.0003 | 4.1 ± 0.4 | 10.4 ± 0.6 | 14.4 ± 3.4 |
| **40°C** | 23.5 ± 0.2 | 2.08 ± 0.22 | 0.34 ± 0.10 | 0.029 ± 0.002 | 0.0048 ± 0.0012 | 0.0015 ± 0.0004 | | | |
| **45°C** | 24.6 ± 0.2 | 1.52 ± 0.20 | 0.19 ± 0.15 | 0.023 ± 0.001 | 0.0021 ± 0.0016 | 0.0008 ± 0.0005 | | | |
| ***Polio*** | | | | | | | | | |
| **25°C** | 17.8 ± 0.3 | 1.76 ± 0.19 | 0.34 ± 0.07 | 0.040 ± 0.005 | 0.0040 ± 0.0008 | 0.0008 ± 0.0016 | | | |
| **30°C** | 23.1 ± 0.2 | 2.20 ± 0.23 | 0.63 ± 0.07 | 0.018 ± 0.002 | 0.0095 ± 0.0018 | 0.0007 ± 0.0002 | 10.7 ± 1.0 | | |
| **35°C** | 29.6 ± 0.3 | 2.96 ± 0.40 | 1.20 ± 0.34 | 0.021 ± 0.003 | 0.0115 ± 0.0042 | 0.0007 ± 0.0002 | | 11.0 ± 1.1 | 23.2 ± 0.5 |
| **40°C** | 45.0 ± 0.6 | 4.50 ± 0.46 | 1.21 ± 0.23 | 0.024 ± 0.003 | 0.0103 ± 0.0022 | 0.0011 ± 0.0002 | | | |
| **45°C** | 52.6 ± 0.5 | 5.26 ± 0.05 | 1.31 ± 0.12 | 0.055 ± 0.003 | 0.0044 ± 0.0006 | 0.0006 ± 0.0001 | | | |
| ***HRV-C*** | | | | | | | | | |
| **25°C** | 9.6 ± 0.1 | 0.71 ± 0.01 | 0.10 ± 0.01 | 0.154 ± 0.004 | 0.0030 ± 0.0004 | 0.0010 ± 0.0004 | | | |
| **30°C** | 12.7 ± 0.1 | 1.09 ± 0.04 | 0.23 ± 0.05 | 0.059 ± 0.003 | 0.0035 ± 0.0020 | 0.0011 ± 0.0003 | 14.3 ± 1.1 | 15.3 ± 1.7 | 34.3 ± 1.8 |
| **35°C** | 23.1 ± 0.3 | 2.31 ± 0.15 | 0.63 ± 0.12 | 0.028 ± 0.002 | 0.0045 ± 0.0010 | 0.0008 ± 0.0002 | | | |
| **40°C** | 28.2 ± 0.3 | 2.28 ± 0.21 | 0.56 ± 0.11 | 0.018 ± 0.002 | 0.0037 ± 0.0008 | 0.0010 ± 0.0002 | | | |
| **45°C** | 42.3 ± 0.3 | 4.23 ± 0.49 | 1.08 ± 0.18 | 0.013 ± 0.001 | 0.0054 ± 0.0011 | 0.0009 ± 0.0002 | | | |

*standard deviation

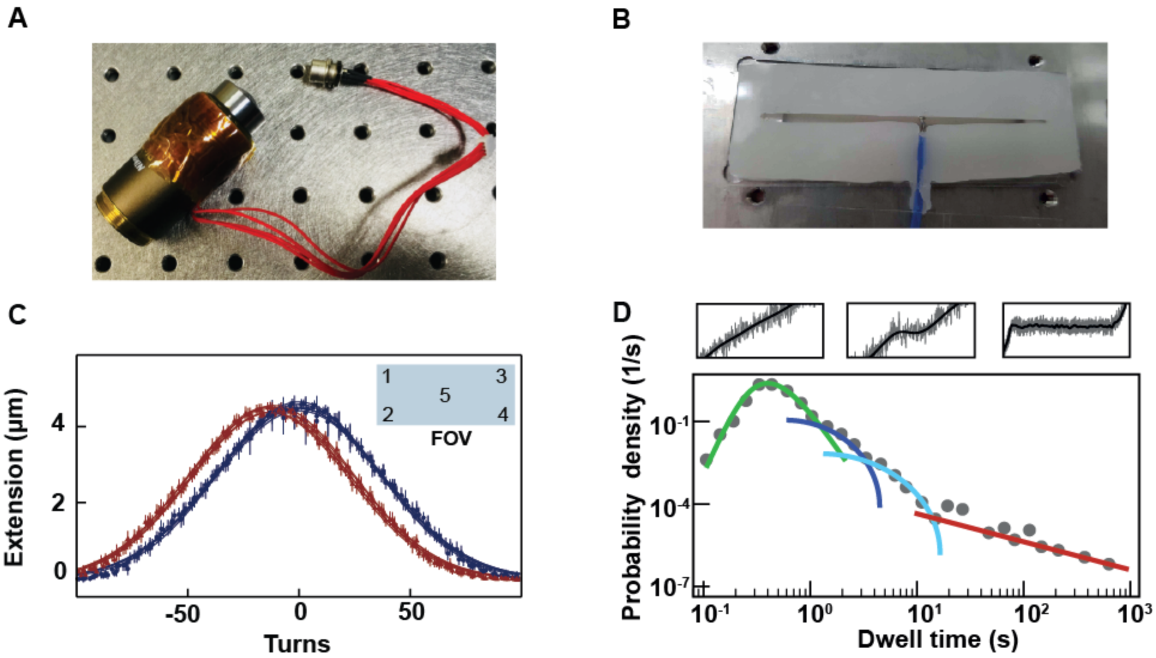


**Supplementary Figure 1: Temperature control system in the high throughput magnetic tweezers assay and dwell times distribution description. (A)** The flexible resistive foil heater (HT10K, Thorlabs) is wrapped around the oil immersion objective. The heating foil is further insulated by several layers of Kapton tape (KAP22-075, Thorlabs). **(B)** Macroscopic temperature probe incorporated into a flow cell. **(C)** Rotation extension curve for a single MyOne magnetic bead tethered by a ~20.6 kbp dsDNA molecule and measured in five different positions of the field of view (see insert) at 25°C (dark blue) and 45°C (red). The extension is measured every 5 turns steps. The error bars denote the standard deviation of the median position of the magnetic bead along the z-axis. **(D)** Dwell times distribution of 45 traces of HRV-C RdRp activity, acquired at 30pN, 35°C and 1mM [NTP]. The distribution is decomposed in four parts: the dwell time <1 s describe nucleotide addition burst without pause, and are described by a gamma distribution (green solid line, fit) for N=10, i.e. the convolution of the exponentially described ten consecutive nucleotide addition cycles; two exponentially pauses, coined Pause 1 (~2 s lifetime, dark blue solid line, MLE fit) and Pause 2 (~10 s lifetime, light blue solid line, MLE fit); pauses longer than ~20 s are well described by a power law, such as $t^{-3/2}$ (red line, MLE fit), which describes a backtracking behavior. The three inserts above the distribution are captures of an RdRp trace which presents the behavior described in the dwell time distribution below, i.e. no pause (gamma distribution), a short pause (exponential distributions) and a long pause (power law distribution). The mathematical expression of the MLE fit is described in the **Material and Methods** section.

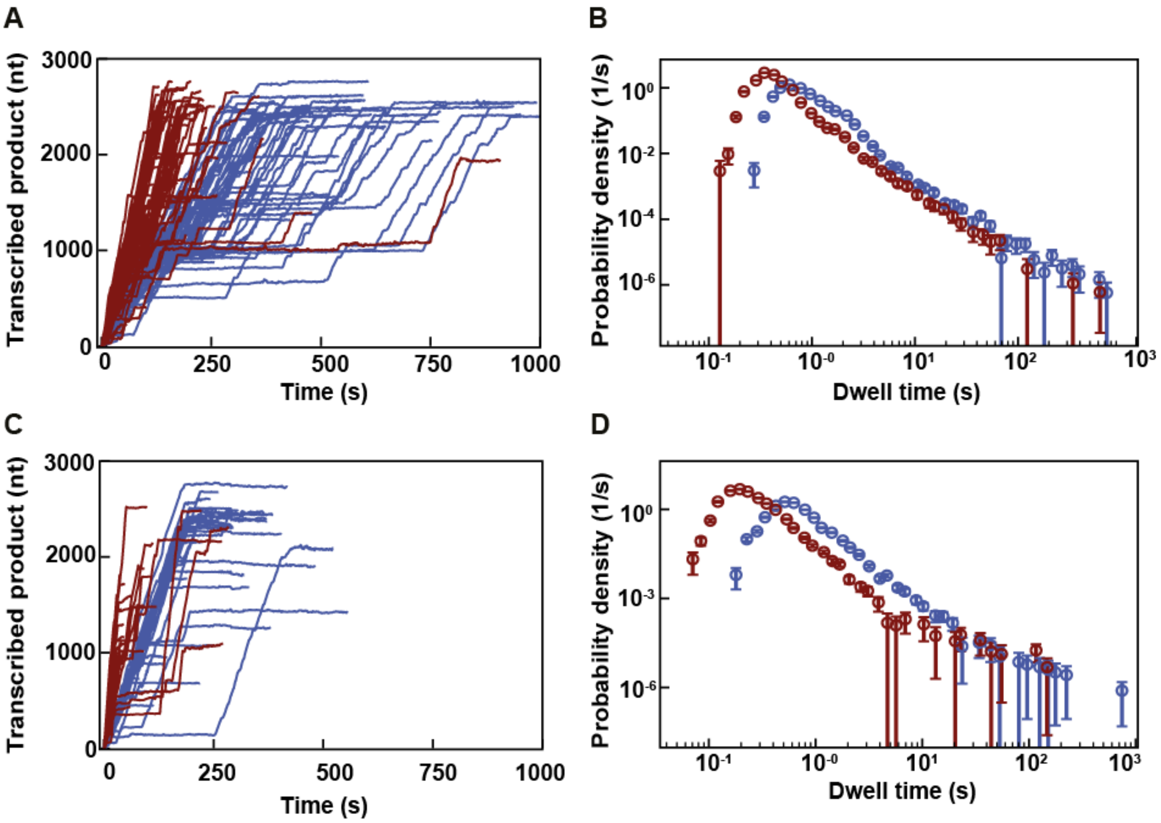


**Supplementary Figure 2: Replication activity of Φ6 and PV RdRps at 25°C (blue) and 45°C (red).** All data were acquired during a single acquisition at 58 Hz and low pass filtered 0.5 Hz. **(A)** Traces of Φ6 RdRp activity at 25°C and 45°C acquired at 30 pN applied force, 1 mM ATP/GTP, 0.2 mM CTP/UTP. **(B)** The corresponding dwell time distributions. **(C)** Traces of PV RdRp activity acquired at 30 pN applied force, 1 mM NTPs and the **(D)** corresponding dwell time distribution.

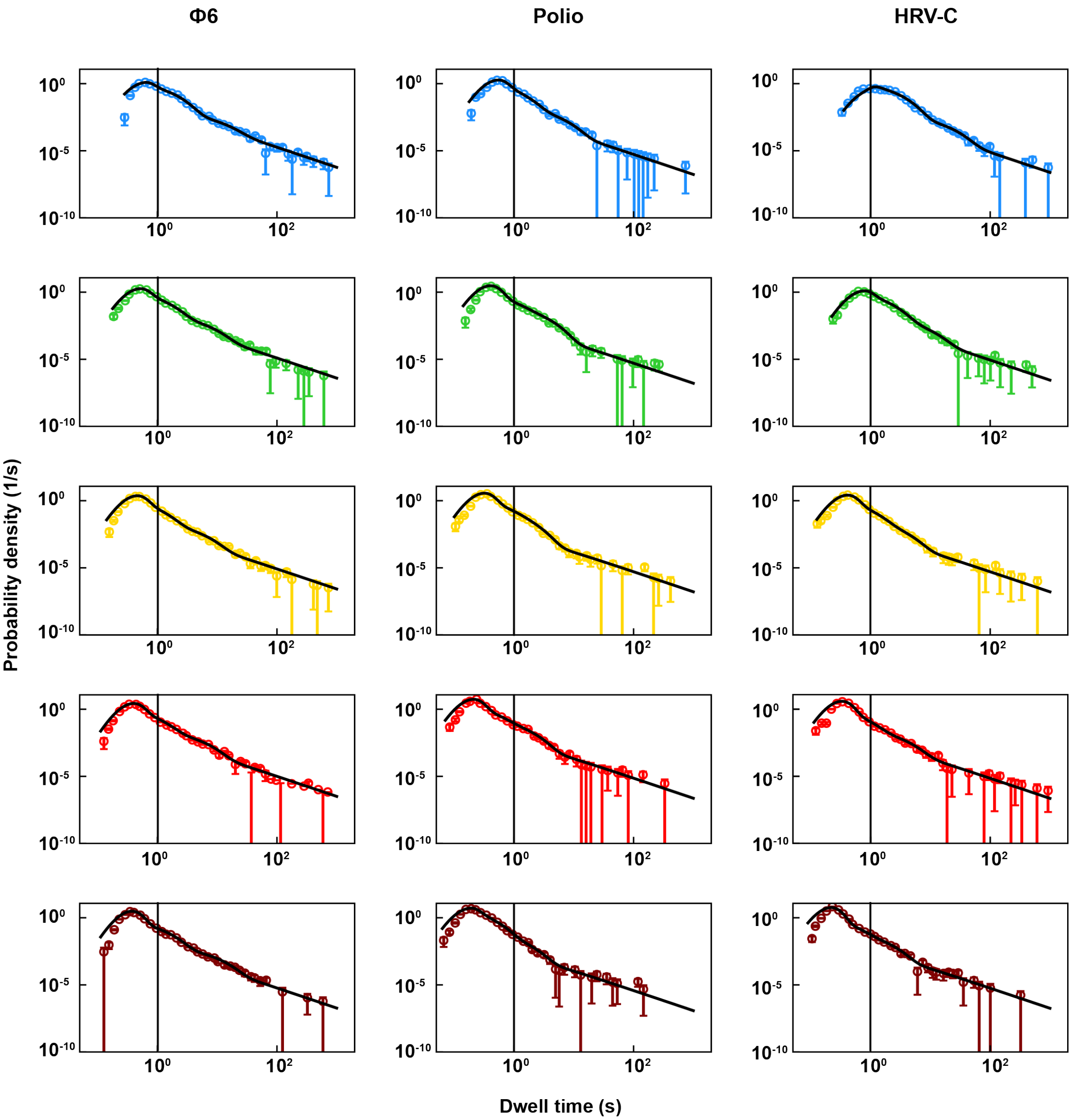


**Supplementary Figure 3: The probability density distributions and MLE fits (solid black lines) of the dwell times extracted from Φ6, poliovirus and HRV-C RdRps' activity traces at different temperatures.** Blue: 25°C, light blue: 30°C, green: 35°C, yellow: 40°C, red: 45°C. A vertical line is added at 1s to illustrate the shift in the gamma peak position. In the HRV-C dwell time distribution at 40°C, the MLE fit is constrained to $k_2 > 0.2\ s^{-1}$. The bins error bars are 36% confidence interval from 1000 bootstraps procedure.


1. Kriegel, F., Matek, C., Drsata, T., Kulenkampff, K., Tschirpke, S., Zacharias, M., Lankas, F. and Lipfert, J. (2018) The temperature dependence of the helical twist of DNA. *Nucleic Acids Res*, **46**, 7998-8009.
